# LOPIT-DC: A simpler approach to high-resolution spatial proteomics

**DOI:** 10.1101/378364

**Authors:** Aikaterini Geladaki, Nina Kočevar Britovšek, Lisa M. Breckels, Tom S. Smith, Claire M. Mulvey, Oliver M. Crook, Laurent Gatto, Kathryn S. Lilley

## Abstract

Hyperplexed Localisation of Organelle Proteins by Isotope Tagging (hyperLOPIT) is a well-established method for studying protein subcellular localisation in complex biological samples. As a simpler alternative we developed a second workflow named Localisation of Organelle Proteins by Isotope Tagging after Differential ultraCentrifugation (LOPIT-DC) which is faster and less resource-intensive. We present the most comprehensive high-resolution mass spectrometry-based human dataset to date and deliver a flexible set of subcellular proteomics protocols for sample preparation and data analysis. For the first time, we methodically compare these two different mass spectrometry-based spatial proteomics methods within the same study and also apply QSep, the first tool that objectively and robustly quantifies subcellular resolution in spatial proteomics data. Using both approaches we highlight suborganellar resolution and isoform-specific subcellular niches as well as the locations of large protein complexes and proteins involved in signalling pathways which play important roles in cancer and metabolism. Finally, we showcase an extensive analysis of the multilocalising proteome identified via both methods.

## Introduction

It is well established that the level of complexity of the human proteome extends far beyond the number of gene products expressed by the genome in a cell^1,2^. Protein subcellular localisation is an important aspect of this complexity since the compartmentalisation of eukaryotic cells and the dynamic distribution of proteins between different organelles are intertwined with the regulation of cellular function^3^. Perturbations in protein subcellular location have serious clinical implications and there are many human diseases caused by abnormal protein expression in combination with aberrant localization^4–9^. Subcellular proteomics studies have led to novel discoveries regarding disease mechanisms, generating new models to link mutations to certain disorders^2,10–24^. Therefore, creating a complete and comprehensive organelle map for each tissue type or cell line under each possible physiological or pathological condition has the potential to significantly benefit drug discovery programs.

Lately, advances in large-scale proteomics technologies^25–27^ have led to spatial proteomics studies which have provided useful insights regarding organelle composition, dynamics and function in health and disease and across a range of different species and cell types^4, 28^. Elaborate subcellular fractionation protocols coupled to differential or density-gradient centrifugation and downstream mass spectrometry-based proteomics methods have been the gold standard for protein subcellular localization analysis for many years and hundreds of subcellular proteomics studies have been published, aiming to characterise all major organelles, macromolecular structures and multiprotein complexes in eukaryotic cells^1–3,16,28–40^. Methods for subcellular fractionation alternative to centrifugation have also been developed and applied to subcellular proteomics research with variable success^1,34,35,41^.

Furthermore, recent advances in the field of quantitative proteomics have greatly contributed to the evolution of spatial proteomics studies. One of the most powerful such strategies has been the development of isotope-labelling methods which allow for the simultaneous analysis of many different biological samples in the same experiment^27, 42^. In *vitro* chemical labelling multiplexing up to 11 different tandem mass tags (TMT) is now possible leading to significant reduction in factors which previously limited subcellular resolution, such as experimental variation arising due to separate mass spectrometry runs and biological sample preparations^42^ as well as missing values resulting from the stochastic nature of mass spectrometry.

Importantly, major developments in bioinformatics including approaches to interrogate spatial proteomics data^43, 44^ and achieve sequence- or annotation-based prediction of protein subcellular localization (reviewed in^37^) have also contributed to the evolution of spatial proteomics methods. Moreover, a large number of publicly-accessible organelle databases and web-based resources have also been developed, some of which link subcellular proteomics data to functional datasets as well as disease relevance and animal model information; two important such resources are UniProt^45^ and The Human Protein Atlas^46^ and some others are reviewed in^37^.

One of the methods that allow for the simultaneous analysis of multiple subcellular structures in complex biological mixtures is termed Localisation of Organelle Proteins by Isotope Tagging (LOPIT). LOPIT was first developed more than a decade ago to globally identify, quantify and assign cellular proteins to their respective subcellular niche^47, 48^ and does not rely on absolute organelle purification (thereby circumventing the problems associated with it) but is based on the measurement of the distributions of cellular proteins across multiple density gradient fractions. In the context of this technique, protein localisation is assigned by comparing the distributions of unlabelled proteins to those of known, well curated organelle markers using quantitative mass spectrometry coupled to multivariate statistical analysis and machine-learning approaches^44^. Finally, LOPIT has been applied for the study of the subcellular proteomes of the HEK293 human kidney cell line, *A. thaliana* roots, D. *melanogaster* embryos, *S. cerevisiae* cells and the DT40 lymphocyte cell line^49–55^.

Recently, an improved version of LOPIT called hyperplexed LOPIT (hyperLOPIT) has been developed and applied to the study of the E14TG2a mouse embryonic stem cell subcellular proteome^56^. The hyperLOPIT protocol integrates novel approaches for sample preparation, mass spectrometry data acquisition and multivariate data analysis to create high resolution protein subcellular localisation datasets. Its application results in subcellular location assignment for thousands of proteins and functional complexes with excellent resolution and enables the novel classification of proteins whose subcellular distribution was previously unknown. It also returns information on proteins that demonstrate intermediate distributions between multiple subcellular compartments, which in turn reflects protein functions involved in important aspects of cell biology. Importantly, hyperLOPIT provides information on a cell-wide scale, unlike proximity tagging methods^57^ designed to identify proteins associated with discrete subcellular niches which only return limited data per experiment and do not readily scale to reveal proteins with multiple locations.

Variations on the hyperLOPIT method have recently been employed by Beltran *et al.*, who integrated a temporal component to the spatial proteomics workflow^58^ to analyse human lung fibroblast cytomegalovirus infection and Jadot *et al.*, who used Nycodenz and sucrose density gradient centrifugation to determine the rat liver organelle proteome^59^. Moreover, by using principles of book-keeping, these authors were able to estimate protein distribution across eight major organelles. A label-free alternative to LOPIT termed Protein Correlation Profiling (PCP) has also been developed and applied to the study of the centrosome^60^ and lipid droplets^61^ as well as to global organelle analyses^62, 63^. This technique has also been used to study the proteasome complexes of *Plasmodium falciparum*^64^ and has been combined with Stable Isotope Labelling with Amino acids in cell Culture (SILAC) to investigate protein-protein interactions with temporal and stoichiometric resolution^65, 66^.

A method based on differential centrifugation alone, called Dynamic Organellar Maps, was recently applied to SILAC-labelled HeLa cells^67^. In the context of this technique cells are processed according to two schemes: a) after removing the nucleus, SILAC-light cells are fractionated into membrane-enriched pellets and b) SILAC-heavy cells are split into nucleus-, organelle- and cytosol-enriched fractions. Each SILAC-light sample is then pooled with a “reference” SILAC-heavy membrane-enriched fraction and analysed on a mass spectrometer to obtain light/heavy ratios from which a protein’s membrane location is inferred. In parallel, all three SILAC-heavy fractions are analysed to obtain information on global protein distribution between the nucleus, cytosol and other organelles. This workflow has since been updated to include a full label-free (LFQ) and a TMT-labelled option^68^ in which no reference organellar fraction is used. In the LFQ option the nucleus is included in the protein membrane distribution analysis and for TMT labelling only the five post-nuclear fractions are used and two replicates labelled with one TMT10plex set.

The approaches described above differ regarding not only laboratory protocols used for lysis, fractionation or quantitation but also data processing and some variations within the bioinformatics analysis pipelines. Such dissimilarities as well as the use of variable subcellular marker lists for the classification of proteins to organelles make different spatial proteomics datasets difficult to compare. In light of these issues, the MetaMass tool developed by Lund-Johansen *et al.* goes some way towards allowing for reliable comparisons between the outputs of different workflows via the use of standardized organelle markers and a k-means clustering approach^43^.

Here, we introduce Localisation of Organelle Proteins by Isotope Tagging after Differential ultraCentrifugation (LOPIT-DC), a novel spatial proteomics pipeline based on differential centrifugation which, unlike previous methods, allows for cell-wide sampling in a single experiment. This workflow requires less starting material than hyperLOPIT and is a simpler and quicker protocol. We compare the protein subcellular localisation maps produced by the two methods using the U-2 OS cell line and also utilise QSep, a recently developed, freely available tool which aims to objectively and robustly quantify subcellular resolution in spatial proteomics data. Importantly, we evaluate the impact of employing a differential centrifugation-based workflow on global, experiment-wide resolution and compare the results side by side with the equivalent hyperLOPIT output, detailing the strengths and weaknesses of each method. Using both approaches, we highlight suborganellar resolution and protein isoform-specific subcellular niches as well as the locations of large protein complexes and proteins involved in signalling pathways which play important roles in cancer and metabolism. We also showcase an extensive analysis of the multilocalising proteomes identified via both methods. In summary, we present the most comprehensive mass spectrometry-based spatial proteomics map of a human cell line to date and deliver a flexible set of subcellular proteomics protocols from sample preparation through to data analysis.

## Results

### Development of the LOPIT-DC method

Aiming to create a simpler alternative to hyperLOPIT, we developed a second mass spectrometry-based technique for the study of protein subcellular localisation which we named Localisation of Organelle Proteins by Isotope Tagging after Differential ultraCentrifugation (LOPIT-DC, Figure 1). For the development of this method we combined the strengths of our hyperLOPIT protocol with elements of other subcellular fractionation methods that employ differential centrifugation^59–67–68^. Seeing how such approaches suffer from low resolution relatively to our hyperLOPIT approach (Figure S9), but also acknowledging that the hyperLOPIT workflow is a relatively expensive and long protocol which requires a large amount of starting material, we modified our method towards the creation of a workflow which would be faster, cheaper and less resource-intensive than hyperLOPIT while retaining the highest subcellular resolution possible. In this article, we present our LOPIT-DC method as a “best-of-both-worlds” scenario aimed at scientists whose questions do not necessarily require obtaining maximum organellar resolution, those who seek to reduce experimentation time in the context of dynamic studies or those who are limited in starting material amount, resources or funding and therefore are after a more economical solution than hyperLOPIT (Table S3).

**Figure 1:**
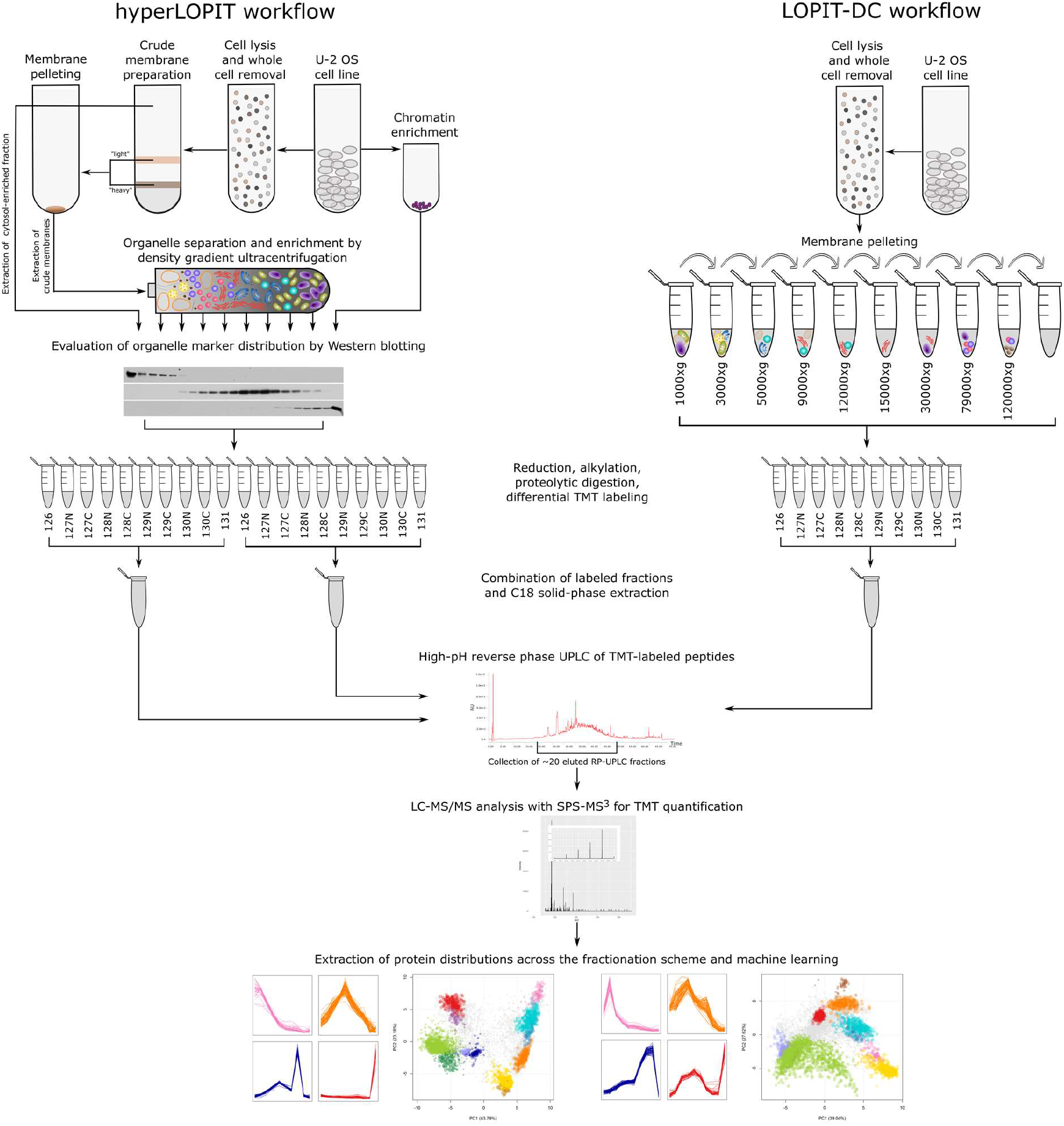
Overview of the hyperLOPIT (left) and LOPIT-DC (right) workflows.

For cell lysis, LOPIT-DC utilises the same approach as hyperLOPIT as it involves a gentle isotonic lysis buffer which keeps organelles as intact as possible while cells are lysed in a ball-bearing cell cracker. This lysis step shows excellent reproducibility and can be optimised for a variety of cell or tissue types. As in hyperLOPIT, cell lysis in LOPIT-DC is followed by a whole cell pre-clearing step which is necessary to remove unlysed cells that could confound downstream analysis. The cell lysis stage in both of our methods is critical as inefficient cell lysis can result in suboptimal organelle recovery which would lead to low protein yield during later steps, reducing overall efficiency. For LOPIT-DC, inefficient lysis would also mean generation of large microsomal particles which would sediment during the initial centrifugation steps. On the other hand, excessive cell lysis can damage sensitive membranes and lead to release of organellar content to the soluble part of the preparation.

The main difference between hyperLOPIT, where the subcellular fractionation part of the protocol is based on density gradient ultracentrifugation, and LOPIT-DC is that fractionation in the latter relies on subsequent ultracentrifugation steps. As a modification of our hyperLOPIT protocol, LOPIT-DC utilises sequential differential centrifugation steps to fractionate the cell lysate into 10 fractions (Table 1). Some of the centrifugation speeds in our LOPIT-DC workflow are similar to those described in^67^ with the addition of 4 extra steps. In contrast to the Dynamic Organellar Maps pipeline, LOPIT-DC is an all-in-one method meaning that all subcellular niches are analysed in a single preparation, thus avoiding any variation arising due to membrane damage and protein leakage.

**Table 1:**
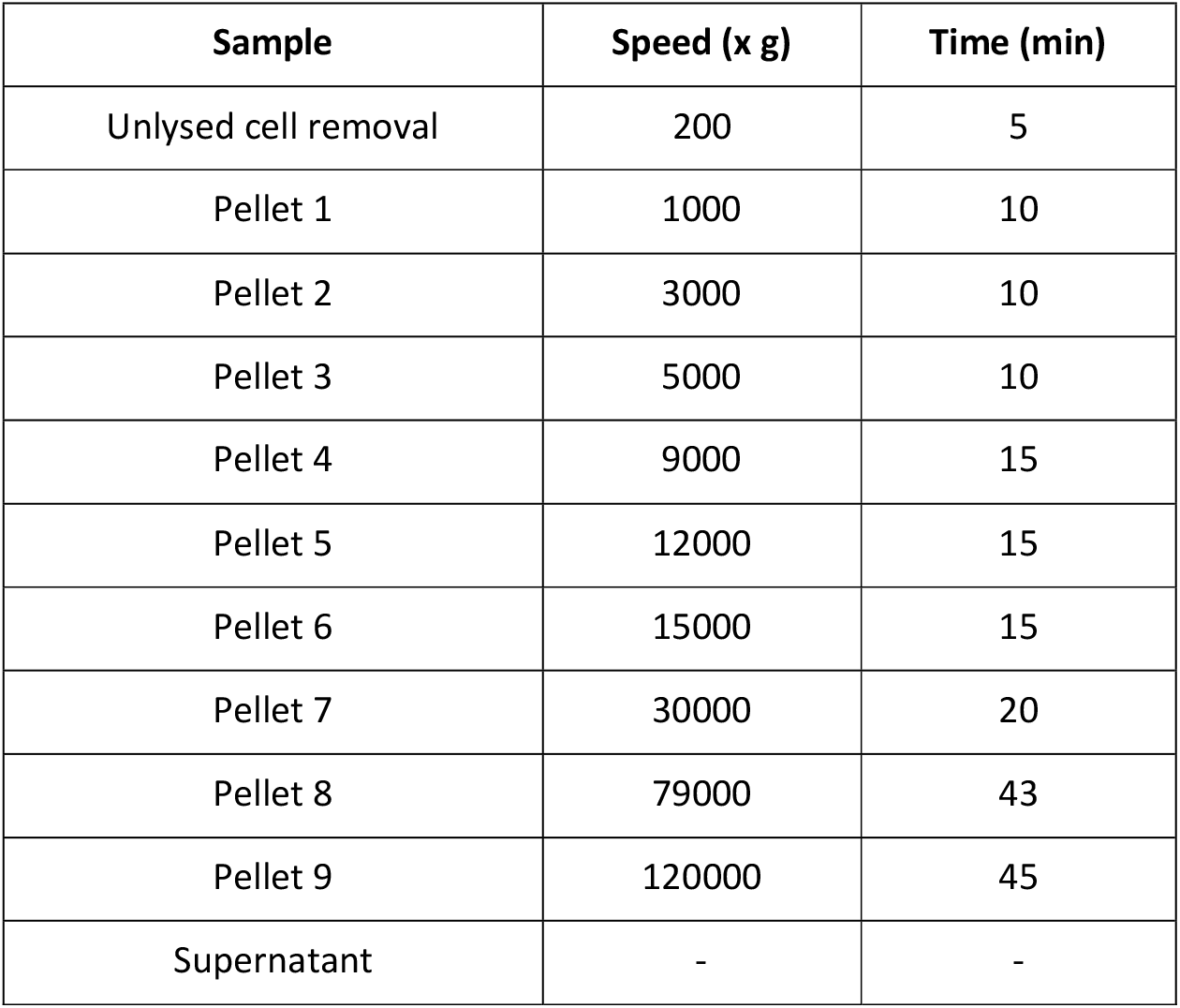
Centrifugation speeds and times for the LOPIT-DC fractionation protocol.

| Sample | Speed (x g) | Time (min) |
| --- | --- | --- |
| Unlysed cell removal | 200 | 5 |
| Pellet 1 | 1000 | 10 |
| Pellet 2 | 3000 | 10 |
| Pellet 3 | 5000 | 10 |
| Pellet 4 | 9000 | 15 |
| Pellet 5 | 12000 | 15 |
| Pellet 6 | 15000 | 15 |
| Pellet 7 | 30000 | 20 |
| Pellet 8 | 79000 | 43 |
| Pellet 9 | 120000 | 45 |
| Supernatant | - | - |

Similar to hyperLOPIT, LOPIT-DC makes use of the TMT multiplexing strategy that allows for simultaneous analysis of all subcellular fractions in a single mass spectrometry run. This approach greatly reduces between-run technical variation and thus the number of missing values per experiment. This is extremely important as presence of a high amount of missing values in such datasets can have a detrimental effect on protein localisation assignment as observed recently by Beltran et al. 2016^58^. Moreover, TMT-based multiplexing allows for a reduction in mass spectrometry time required and thus in cost. As with hyperLOPIT, mass spectrometric analysis in the context of the LOPIT-DC protocol is carried out using the powerful SPS-based MS^3^ technology on the Orbitrap Fusion Lumos Tribrid system. This way quantitative accuracy and precision are maximised, leading to improved spatial resolution (A. Christoforou *et al.*, 2016 and Figure 2 therein).

**Figure 2:**
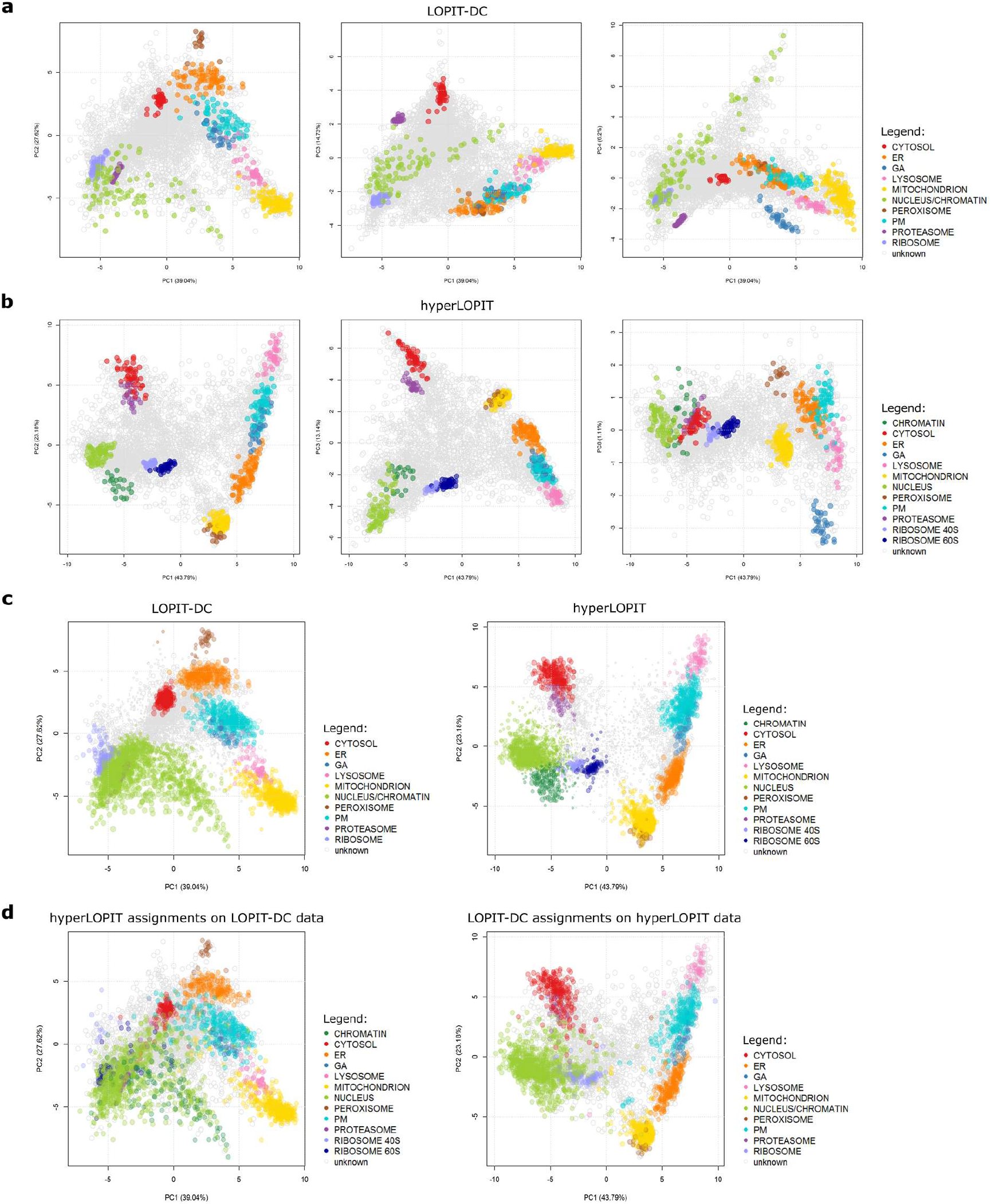
PCA plots of the LOPIT-DC and hyperLOPIT datasets. a) PCA plots of the merged LOPIT-DC dataset overlaid with markers and plotted in different dimensions that display resolution of all organellar clusters; b) PCA plots of the merged hyperLOPIT dataset overlaid with markers and plotted in different dimensions that display resolution of all organellar clusters; c) PCA plots displaying the SVM classification results after applying a 5% FDR cutoff for the LOPIT-DC and hyperLOPIT datasets; d) hyperLOPIT assignments plotted upon the LOPIT-DC data and *vice versa.*

For downstream statistical analysis, LOPIT-DC employs the robust spatial proteomics data analysis workflow based on the freely available MSnbase and pRoloc Bioconductor packages, created using the R statistical programming environment. The workflow^69^ offers simple features such as data import, export, processing and quality control, subcellular marker definition, visualisation and interactive exploration as well as advanced functions such as clustering, protein subcellular localisation classification^70^, novelty detection using semi-supervised learning^51^ and transfer learning^71^. In addition, the pRolocdata package^44^ provides a variety of readily available, annotated and preformatted datasets generated using LOPIT or hyperLOPIT over the years and originating from different species. These open-source, open-development R packages constitute our full data analysis pipeline. They are interactive, user friendly and are accompanied by working examples, full documentation, tutorials and videos allowing users to follow this analysis workflow step-by-step^69^. Moreover, new tools and methods are continuously being integrated to keep at the forefront of advances in machine learning.

### Application of LOPIT-DC on the U-2 OS cell line

LOPIT-DC was applied to U-2 OS cells. This cell line was chosen as it is a well characterised model that has been used for a variety of research purposes. More importantly, a large amount of immunofluorescence-based protein subcellular localisation data obtained using this cell line is publicly available as part of the Cell Atlas project^72^. Therefore, this database is an excellent source of information for the validation of our mass spectrometry-based spatial proteomics observations and has served as such in the past (more details in next section).

We performed three biological replicates of LOPIT-DC on the U-2 OS cell line using on average 70×10^6^ cells per replicate (Figures S1a, S2a, S3a, S4). This way we obtained at least 60 μg of protein in each fraction with P6 being consistently the lowest-yield sample (Figure S3a). LC-SPS-MS^3^ analysis of the U-2 OS LOPIT-DC fractions resulted in identification of 9386 protein groups after replicate merging and, following initial processing and missing value removal, 6837 protein groups with a full reporter ion series remained (Table S1).

Principal Component Analysis (PCA) revealed underlying data structure and the quality and identity of these clusters were further explored by overlaying a collection of manually curated organelle markers on this dataset. As Figures 2a, 3a and 3b show, LOPIT-DC offers superb resolution concerning most major subcellular niches. In more detail, our LOPIT-DC experiments were able to resolve the following 10 major organelle clusters: cytosol, nucleus/chromatin, mitochondrion, peroxisome, lysosome, endoplasmic reticulum (ER), plasma membrane (PM), Golgi apparatus (GA), ribosomes and proteasome. In the merged dataset the chromatin is only partially resolved from the non-chromatin nuclear compartment and the two ribosomal subunits are not separated (Figure 2d, Figure S6). Moreover, the various organelle clusters seem to be organised into three larger groups separated from each other by greater distances: the first group contains the membranous organelles excluding the nucleus, the second group includes the nucleus/chromatin and ribosomes and the third group consists of the cytosolic cluster and proteasome. Importantly, subcellular niches that seem to overlap in principal components 1 and 2 are separated in other dimensions. For example, the GA and PM exhibit overlapping distributions in PCs 1 and 2 but are separated along dimensions 1 and 4. Similarly, the nucleus/chromatin, ribosome and proteasome clusters seem to overlap in dimensions 1 and 2 but these structures are separated from each other along principal components 1 and 3. LOPIT-DC also offers good reproducibility between replicates based on protein yield per fraction (Figure S3a) and the fact that all three of our U-2 OS LOPIT-DC experiments exhibit similar subcellular resolution (Figure S1a, S2a, S4).

**Figure 3:**
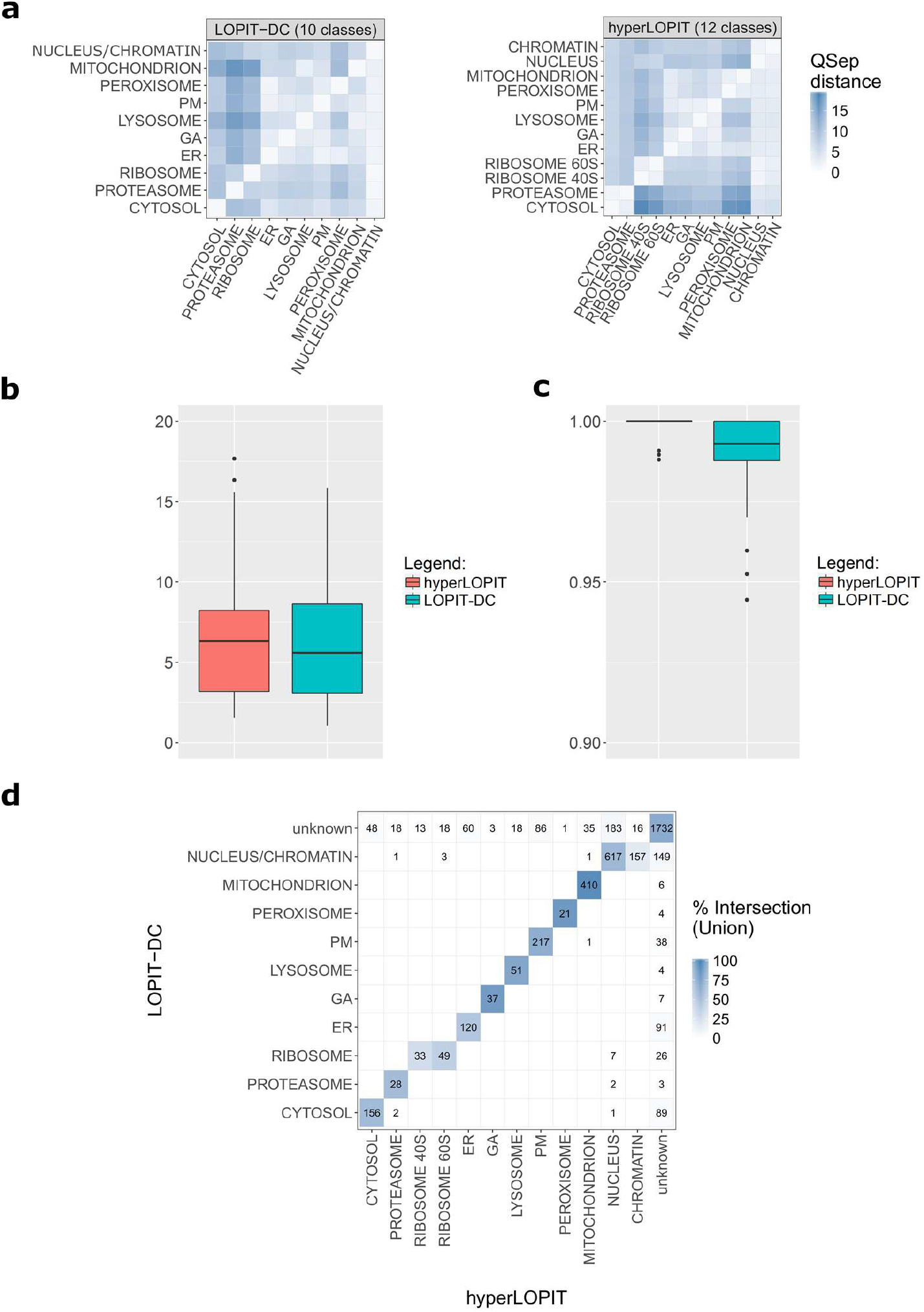
Quantifying the resolution, marker behavior and protein subcellular location assignments for the LOPIT-DC and hyperLOPIT experiments. a) Annotated heatmaps depicting all normalised pairwise distances for the LOPIT-DC and hyperLOPIT datasets; b) Boxplots displaying average normalised pairwise distances for the LOPIT-DC and hyperLOPIT datasets; c) Macro F1 scores for the LOPIT-DC and hyperLOPIT datasets; d) Heatmaps displaying the overlap between the LOPIT-DC and hyperLOPIT protein subcellular localisation assignments (including markers), where the LOPIT-DC dataset is classified using 10 marker classes and the colour code is based on the percentage of intersection (i.e., the number of intersecting proteins is divided by the total number of proteins assigned to that organelle in the LOPIT-DC and the hyperLOPIT data).

Finally, we applied supervised machine learning using a SVM-based classifier in order to predict the subcellular localisation of the unlabelled proteins in the merged U-2 OS LOPIT-DC dataset. We performed classification using 12 (cytosol, nucleus, chromatin, mitochondrion, peroxisome, lysosome, ER, PM, GA, ribosome 40S, ribosome 60S and proteasome) or 10 organelle classes, where the nucleus and chromatin were merged into one cluster and the same was done for the two ribosomal subunits. In both cases, after classification the majority of proteins were assigned to the nuclear, cytosolic and mitochondrial clusters whereas the least populated niches were the proteasome, peroxisome, GA and lysosome. Based on information available in UniProt and the literature we manually set SVM classification thresholds for each subcellular niche allowing for a 5% false discovery rate. This way, approximately 35% of the merged dataset proteins were assigned to their respective subcellular location (Table S2, Figure 2c). Finally, as demonstrated by Table S2 and Figure S6, as expected, using 10 organelle classes rather than 12 improved classification numbers and quality regarding subcellular niches not optimally resolved in LOPIT-DC, such as the ribosomes.

In hyperLOPIT, an additional nuclear chromatin preparation aids in separating the nucleus and chromatin clusters. To investigate if the same is true for our differential centrifugation-based workflow, we prepared and added this chromatin-enriched fraction to our LOPIT-DC analysis. As can be observed in Figure S7, this addition did not significantly improve the resolution of our U-2 OS LOPIT-DC dataset but in turn led to a slight decrease in overall subcellular resolution (Figure S7b), therefore we excluded this chromatin-enriched fraction from our downstream data analysis (see Figures S7b and S7c for details).

### Acquisition of a complete U-2 OS hyperLOPIT dataset

In continuation to the two-replicate dataset we presented as part of Thul et al. 2017^72^, we performed a third replicate of our hyperLOPIT method using U-2 OS cells; here we present this complete dataset and compare it to our LOPIT-DC findings for the same cell line. Importantly, our U-2 OS hyperLOPIT dataset is the largest human (hyper)LOPIT dataset reported thus far. Additionally, this dataset is the most highly resolved mass spectrometry-based human subcellular map published to date.

Our three U-2 OS hyperLOPIT experiments required on average 280×10^6^ cells each. This way we consistently obtained at least 70 μg of protein in each fraction with the exception of the first 5-7 fractions which were pooled for further analysis (Figure S3b). While the LOPIT-DC workflow focuses on simplicity and speed, our hyperLOPIT approach aims at achieving the maximum overall resolution possible leading us to pursue a different TMT labelling strategy. In more detail, we included all density gradient fractions, together with the cytosol- and chromatin-enriched samples, in our hyperLOPIT analysis, ending up with 20 TMT channels per replicate and 60 TMT channels for our merged dataset as opposed to our 10-channel LOPIT-DC dataset and previous hyperLOPIT reports^56^ (Figure 1, Supplementary quantitation table). Due to the large number of samples analysed during our hyperLOPIT experiments the amount of missing values which arose throughout the analysis was higher compared to the LOPIT-DC dataset, leading to the final combined hyperLOPIT dataset being smaller than the combined LOPIT-DC one; following quantitative LC-SPS-MS^3^ analysis of all three of our hyperLOPIT replicates we identified 9558 protein groups which were reduced to 4883 after filtering and concatenating replicates (Table S1). Three of the 60 TMT channels present in our final hyperLOPIT dataset possessed extremely low ion intensity profiles so they were excluded from downstream data analysis to minimise background noise to the data.

As shown in Figure 2b, 12 major subcellular niches were successfully resolved during our hyperLOPIT experiments: the cytosol, nucleus, chromatin, mitochondrion, peroxisome, lysosome, ER, PM, GA, ribosomal subunit 40S, ribosomal subunit 60S and proteasome. Unlike our LOPIT-DC observations, our hyperLOPIT experiments accomplished separation between the two ribosomal subunits as well as the nucleus and nuclear chromatin (Figures 2b, S2b). Furthermore, similarly to the LOPIT-DC data, the organelle classes present in the hyperLOPIT dataset seem to be arranged in four larger groups: the first group contains the cytosol and proteasome, the second the nucleus, nuclear chromatin and ribosomal subunits, the third the mitochondrion and peroxisome and the fourth the membranous organelles of the secretory pathway (lysosome, PM, ER, GA). Notably, while in dimensions 1 and 2 the GA and peroxisome seem to partially overlap with the PM and mitochondrion, respectively, these organelles become entirely separated from each other along principal components 1 and 8. Our three hyperLOPIT replicate experiments also exhibit excellent reproducibility and, importantly, complementary subcellular resolution (Figure S1b, S2b, S3b, S4).

The results from a SVM classification on the hyperLOPIT dataset showed that the majority of unlabelled proteins were assigned to the nucleus (765 proteins), mitochondrion (426 proteins) and PM (267 proteins). A smaller number of proteins was assigned to the GA (7 proteins), peroxisome (9 proteins), proteasome (23 proteins) and ribosomes (23 proteins for the ribosome 40S and 28 proteins for the ribosome 60S), akin to the LOPIT-DC classification results. We found that approximately 42% of the proteins in our hyperLOPIT data were classified as part of a single subcellular niche (Table S2, Figure 2c).

### QSep quantifies the subcellular resolution of spatial proteomics experiments

In order to apply a quantitative metric to our observations concerning the overall resolution of our experiments at the organelle marker level we utilised QSep, a tool which aims to objectively and robustly quantify subcellular resolution in spatial proteomics data and is freely available as part of the pRoloc package. For a detailed description of this function the reader is referred to the Materials and Methods section and^73^.

The QSep metric was applied to the LOPIT-DC (10 organelle classes) and hyperLOPIT (12 organelle classes) datasets in order to quantitatively assess the levels of subcellular resolution our two methods accomplished. The quantitative cluster separation heatmaps in Figure 3a demonstrate that both datasets feature exceptional subcellular diversity and thus spatial resolution, as both heatmaps contain similar colour patterns with a majority of average (light blue) and large (dark blue) normalised pairwise distances across all subcellular clusters. The higher overall resolution afforded by hyperLOPIT is supported by the average normalised pairwise distances describing each dataset, shown in Figure 3b. According to this figure hyperLOPIT displays the highest global, experiment-wide subcellular resolution.

Within the LOPIT-DC dataset the smallest normalised distances correspond to the peroxisome/ER, ER/PM and lysosome/mitochondria pairs (Figure 3a, left); this is in concordance with our PCA plot observations according to which these subcellular niches are positioned close together in PCA space, forming a continuum of clusters. Smaller distances in the heatmap are attributable to differences in cluster size: for example, the nucleus is a much larger cluster than the ribosome and so the average normalised distance between the two is greater than the distance within the nuclear cluster. However, when examining in reverse, the distance between the ribosomes and nucleus is 6.06 times greater than the within-ribosome distance. On the other hand, the largest normalised distances in this dataset are those between various organelles and the proteasome or ribosome. This is expected, as both the ribosome and proteasome are clearly very well separated from the membrane-bound organelles of the secretory pathway as well as the mitochondrion along dimensions 1 and 2.

Within the hyperLOPIT dataset the smallest normalised pairwise distances are those between the two ribosomal subunits and the nucleus as well as between the cytosol and the proteasome (Figure 3a, right). In the case of the ribosome 40S/nucleus and ribosome 60S/nucleus pairs this result is due to differences in cluster size, as also observed in the case of the LOPIT-DC data. Concerning the cytosol/proteasome pair, the low distance value reflects the fact that these two clusters partially overlap in principal components 1 and 2; another example of small normalised distances due to overlapping clusters along these dimensions is the case of the PM/GA pair. On the contrary, the largest normalised distances in the hyperLOPIT dataset correspond to various organelles paired with the cytosol. This demonstrates that the cytosol exhibits the best separation from the rest of the organelle clusters in our hyperLOPIT data.

We next employed QSep in order to assess the overall subcellular resolution of our LOPIT-DC and hyperLOPIT data compared to a variety of publicly available spatial proteomics datasets. In the context of this analysis we applied minimal data post-processing and used, whenever possible, the annotation provided by the original publications. Moreover, we only considered organelle classes defined by at least 7 subcellular markers. In the cases where multiple replicates of a dataset were available, we used the combined dataset as opposed to individual replicate experiments and, in the cases where a combined dataset was unavailable, we used just the first (or only) replicate experiment provided by the authors. Furthermore, any missing values present in the datasets were retained during our QSep analysis. Figure S9 displays the distributions of the global average normalised distances stemming from all subcellular clusters for each dataset, with the datasets ordered according to experiment-wide median between-cluster distance. Interestingly, as demonstrated by this figure, our LOPIT-DC and hyperLOPIT data exhibit the best overall, experiment-wide resolution compared to all the other datasets, with the U-2 OS hyperLOPIT data being top-ranked. This result demonstrates the superb quality of both of our datasets. A detailed description of the datasets used for this analysis as well as a more extensive comparison using additional datasets are presented in^73^.

### Macro F1 scores and SVM-based classification results demonstrate similarity between datasets generated using LOPIT-DC or hyperLOPIT

After exploring the similarities and differences between our LOPIT-DC and hyperLOPIT datasets regarding subcellular resolution at the marker level, we expanded our characterisation to the level of protein subcellular localisation prediction. As mentioned above, in order to assign the unlabelled proteins in our data to a unique subcellular location we performed SVM-based supervised machine learning using 10 organelle classes for the LOPIT-DC dataset and 12 for the hyperLOPIT data. As a first step in assessing classifier performance we examined the macro F1 scores^69^ (harmonic mean of precision and recall) obtained after SVM parameter optimisation for each of our datasets. Macro F1 score values range from 0 to 1 and a high score suggests that the marker proteins in the test dataset are consistently assigned to the correct subcellular location by the algorithm^69^. As shown in Figure 3c, the average F1 scores acquired during SVM parameter optimisation using our core marker set were optimal for both of the datasets as both values were very close to 1. At the level of individual organelle scores the classifier performed best for the hyperLOPIT dataset (Figure S5b) and slightly worse for the LOPIT-DC data, specifically in the cases of the lysosome and PM (Figure S5a).

Figure 2c shows the U-2 OS LOPIT-DC (left) and hyperLOPIT (right) datasets after SVM-based protein subcellular location classification followed by 5% FDR filtering. A larger number of proteins was assigned to a unique location in the LOPIT-DC data compared to our hyperLOPIT dataset but the proportion of classified proteins was slightly higher in the hyperLOPIT dataset (42%) as opposed to the LOPIT-DC data (35%) (Table S2, Figure 2). We proceeded to a comparison between the SVM predictions obtained for each dataset in the form of contingency matrices and heatmaps, aiming to visualise and explore the level of agreement achieved by our two distinct workflows and the potential emergence of method-specific biases towards particular organelles. Strikingly, our two datasets exhibit an outstanding level of agreement as the majority of the proteins which were assigned to a unique subcellular compartment in one dataset were classified to the same location in the second dataset (Figure 3d). Furthermore, the vast majority of the “chromatin” and “nucleus” as well as the “ribosome 40S” and “ribosome 60S” hyperLOPIT classifications were assigned to the “nucleus/chromatin” and “ribosome” niches by LOPIT-DC, respectively. Importantly, the very few mismatches which can be identified between the LOPIT-DC and hyperLOPIT organelle assignments are either false positives resulting from the 5% FDR filtering process or proteins that could be labelled as residents of either of the predicted locations according to published evidence. Furthermore, it is apparent from Figure 3d that the majority of classification disparities between the LOPIT-DC and hyperLOPIT data stem from cases where a protein was assigned to a unique subcellular niche in one dataset but remained unlabelled in the second dataset. In this case, the highest number of proteins which were labelled as “unknown” in the LOPIT-DC dataset was assigned to the nucleus in the hyperLOPIT dataset and vice versa. Finally, the heatmap presented in Figure 3d is color-coded according to the percentage of intersection between the LOPIT-DC and hyperLOPIT SVM-based organelle assignments; the intersection in this case is calculated by dividing the number of matching LOPIT-DC and hyperLOPIT classifications by the union of the LOPIT-DC and hyperLOPIT assignments for that same organelle. Additional comparisons such as the ones including missing proteins or comparing the results of SVM-based protein subcellular localisation classification using 12 instead of 10 organelle classes for the LOPIT-DC dataset are presented in Figure S8.

If we take the hyperLOPIT dataset, visualise it by PCA and then annotate it with the localisations determined by the LOPIT-DC method (Figure 2d, right) we see that these localisations (as highlighted by coloured points) form similar clusters to what we observe in the original hyperLOPIT plot (Figure 2c, right). The main difference is that the “nucleus” and “chromatin” hyperLOPIT classes from the original hyperLOPIT experiment now correspond to one “nucleus/chromatin” class and, similarly, the “ribosome 40S” and “ribosome 60S” classes correspond to a single “ribosome” group as dictated by the 10-class LOPIT-DC classifications. Similarly, if we take the original LOPIT-DC dataset, visualise it by PCA and annotate it with the localisations determined by the hyperLOPIT data (Figure 2d, left) we find that the localisations form similar clusters to those found in the original LOPIT-DC dataset (Figure 2c, left). Interestingly, the distributions of the two hyperLOPIT subnuclear class assignments in this plot indicate that our LOPIT-DC experiments achieved at least partial separation between the chromatin and nucleus clusters without the need for a separate chromatin enrichment step. In conclusion, the above observations demonstrate that the SVM-based protein subcellular localisation classifications acquired for our LOPIT-DC and hyperLOPIT data are transferable between the two datasets, indicating their extremely high agreement.

### Transfer learning showcases the benefit of combining different approaches to tackle protein subcellular localisation prediction

As described in^71^, transfer learning can be used for the meaningful integration of heterogeneous data sources in order to improve overall protein subcellular location classification given an optimal combination of the datasets provided. Our transfer learning approach is based on the integration of a primary experimental spatial proteomics dataset and an auxiliary dataset and, as we have previously demonstrated, results in the assignment of proteins to their respective subcellular niche with higher generalisation accuracy than standard supervised machine learning workflows using a single information source^71^. The aim behind implementing such an approach is to support and complement the primary data with secondary annotation features without compromising their integrity, with the user possessing complete control over the amount of auxiliary data to incorporate into the learning process.

The first step during transfer learning is free parameter optimisation for the classifier. Our transfer learning approach uses a *k*-NN classifier and requires optimisation of two different sets of parameters: the first set is the *k*’s necessary for the nearest neighbour calculations for the primary and auxiliary datasets and the second is the organelle class weights, one per class, which determine the proportion of primary and secondary data to be used for learning and range between 0 and 1. A weight of 1 implies that all weight is given to the primary data source, meaning that the final result relies exclusively on the primary experimental dataset and ignores the auxiliary data source provided. Conversely, a weight of 0 indicates that all weight is given to the auxiliary data, representing a situation where the primary source of information is completely ignored and only the secondary dataset is considered. A weight of 0.5 implies that both data sources are equally used during learning and so contribute equally to the final result. The optimal combination of subcellular class-specific weights for a given primary and auxiliary data pair is identified during the algorithm’s optimisation task and the results can be directly plotted as a bubble plot illustrating the proportion of best weights observed during the optimisation phase for each organelle class.

We have previously shown that transfer learning is particularly useful for organelle classes which are not optimally resolved in the primary experimental data. Given that and in order to explore whether we can improve protein subcellular localisation assignment and thus gain new information by making use of the unique features and strengths of each of our two subcellular fractionation methods, we next proceeded to apply transfer learning on the LOPIT-DC and hyperLOPIT U-2 OS cell datasets. Since our prior analysis, described above, indicated that hyperLOPIT achieved higher overall resolution than LOPIT-DC during our experiments and aiming to maximise subcellular resolution after classification, we used the hyperLOPIT data as the primary information source and the LOPIT-DC dataset as the auxiliary data. Figure 4a shows the distribution of the class-specific weights selected over 100 test partitions of the transfer learning algorithm applied to the two datasets. As evident in this figure, the weight distributions corresponding to each dataset closely reflect the resolution achieved by either hyperLOPIT or LOPIT-DC during our experiments. In more detail, the distribution of the best identified weights is skewed towards 1 for just under half of subcellular compartments suggesting that the proportion of neighbours to use during protein subcellular location classification to these organelles should be predominantly primary and indicating their better resolution in hyperLOPIT. However, this is not true for all subcellular niches: the cytosol was assigned best weight of 0 in 78% of runs, signifying that auxiliary data should be used to classify to it. This observation reflects the overlapping distributions exhibited by the cytosol and proteasome in the hyperLOPIT dataset and, in turn, their superb separation from each other and all other organelles in the LOPIT-DC data. Furthermore, half of the subcellular compartments were assigned weights of 0.5 indicating that each dataset should contribute equally to the classification of those subcellular organelles. Finally, the macro F1 scores obtained after weight optimisation and classification of our unannotated proteins demonstrate that including the auxiliary data in the classification leads to an increase in classifier prediction relative to the generalisation accuracy acquired using the hyperLOPIT dataset alone (Figure 4b). Importantly, our findings highlight the merit of integrating our two spatial proteomics methods in order to achieve optimal classification of proteins to organelles.

**Figure 4:**
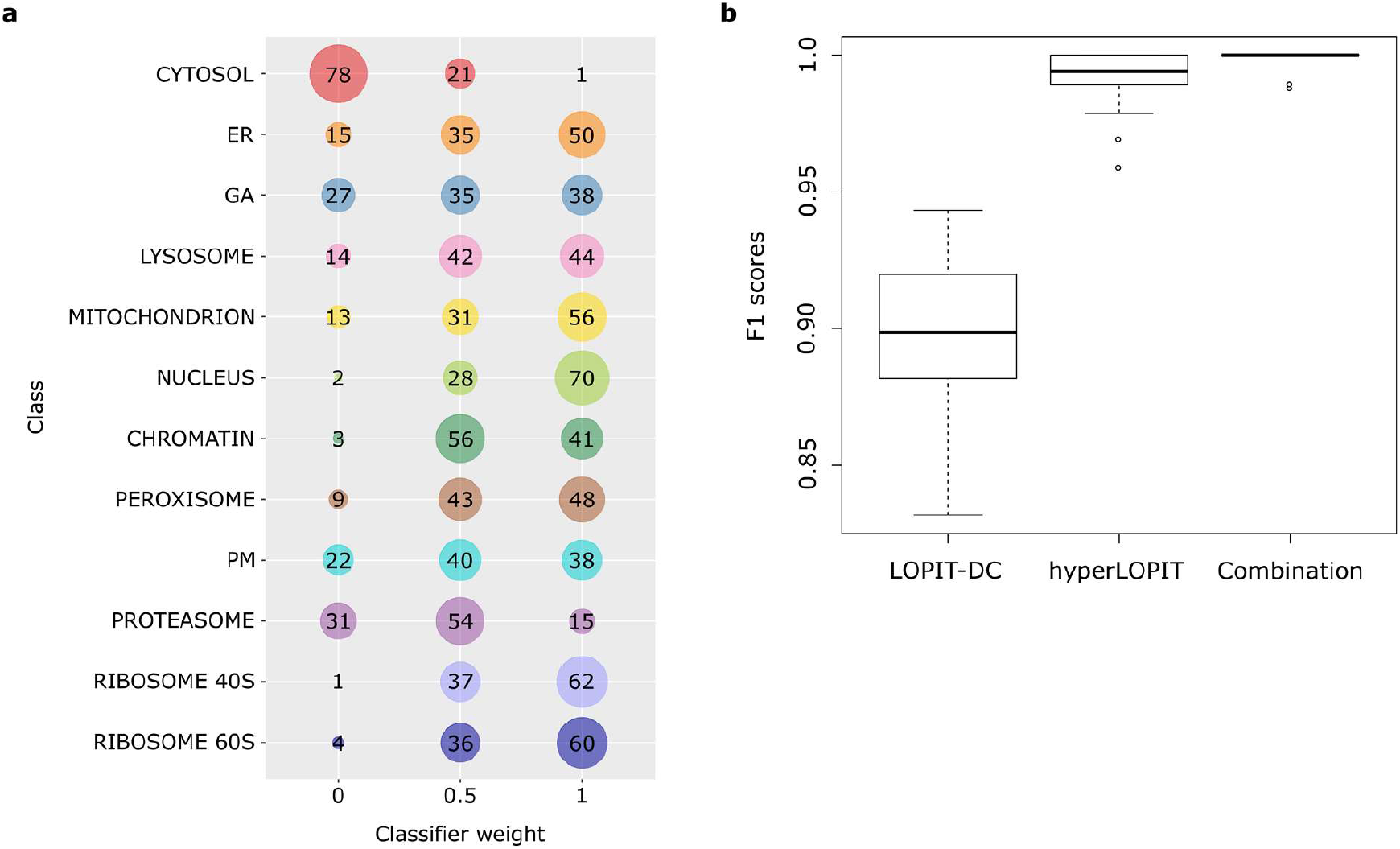
Transfer learning using the hyperLOPIT and LOPIT-DC datasets as the main and auxiliary data sources, respectively. a) Visualisation of the transfer learning parameter optimisation step: each row shows the frequency of observed weights, along the columns, for a specific organelle class, with large circles representing higher observation frequencies; b) PCA plot of the hyperLOPIT dataset after classification, with point size being proportional to classification score; c) F1 scores of the LOPIT-DC dataset, the hyperLOPIT datasets and the combination of the two.

### Proteins in transit account for the largest proportion of the proteins identified by both methods

Apart from the proteins unambiguously classified to a unique subcellular niche, more than half of the proteins in each of our datasets remained unlabelled after SVM-based subcellular location prediction and subsequent 5% FDR filtering. More specifically, 65% and 58% of the total number of proteins initially present in our analysis remained unclassified in the LOPIT-DC and hyperLOPIT data, respectively. These proteins might: 1) reside in more than one subcellular compartments, 2) associate with dynamic components, 3) be active traffickers between different organelles or/and 4) belong to subcellular structures for which no known markers were included in the analysis. Importantly, these unassigned proteins constitute an important part of any spatial proteomics experiment as they most likely represent the dynamic effectors of protein (re)localisation-dependent changes within the cell. Furthermore, many of these multilocalising proteins are also multifunctional with protein function in these cases often depending on a specific subcellular niche and it is such potential translocators which have in many instances been identified as responsible for causing disease in cases of aberrant protein trafficking or/and aggregation^4–6,8^. In many of these cases, early stages of disease can be identified by protein translocation events which precede changes in gene expression. These changes often do not result in overall protein abundance alterations and can therefore only be studied at the subcellular level. Examples of multilocalising proteins include, among others, signalling molecules, transporters, cytoskeletal components, transcription factors, proteins associated with vesicles or junctions, secreted factors and moonlighting proteins. Due to the significance of the diverse roles these molecules play in the cell we proceeded to further explorative analysis of the proteins which were labelled as “unknown” in both of our datasets.

Initially, we utilised the immunofluorescence-based, U-2 OS cell-specific protein subcellular localisation information available as part of the Cell Atlas database in order to investigate the subcellular distribution of our unlabelled proteins. As illustrated in Figures 5a and 5b, the location which harbors the majority (300/200+ occurrences) of the proteins which remained unclassified in both the LOPIT-DC and hyperLOPIT datasets is the cytosol according to the Cell Atlas U-2 OS data. This is expected as many known translocators are soluble cytosolic proteins capable of migrating towards different organelles to exert their function(s). The rest of the unassigned proteins in both of our datasets are mostly (50+ occurrences) distributed to the nucleoplasm or vesicles or are shared between the nucleoplasm and cytosol or cytosol and plasma membrane based on the Cell Atlas database. Strikingly, these distribution patterns reflect the fluid nature of the above compartments: transcriptional regulators and many other kinds of proteins constantly travel between the nucleoplasm and the cytosol; proteins traffic to all organelles as well as the extracellular space and are also led to the degradation pathway via many types of vesicles; the plasma membrane, being the site of secretion and endocytosis as well as intercellular communication, possesses an exceptionally dynamic protein composition. Interestingly, there are obvious differences between the distribution patterns of the proteins which were identified as “unknown” as part of our LOPIT-DC or hyperLOPIT data across the locations defined by the Cell Atlas. In more detail, the subcellular niche that contains the second largest number of unlabelled proteins in the case of the LOPIT-DC dataset is the nucleoplasm, followed by, in order, the nucleoplasm/cytosol combination, vesicles and cytosol/plasma membrane combination. On the other hand, the location harboring the second highest amount of unclassified proteins regarding the hyperLOPIT data is the vesicles, followed by the nucleoplasm as well as the nucleoplasm/cytosol and cytosol/plasma membrane combinations. This discrepancy might indicate that, while both methods primarily identify translocators associated with the cytosol, hyperLOPIT is able to capture the vesicle-associated dynamic proteome more effectively than LOPIT-DC which in turn most efficiently covers the nucleus-associated multilocalising proteome.

**Figure 5:**
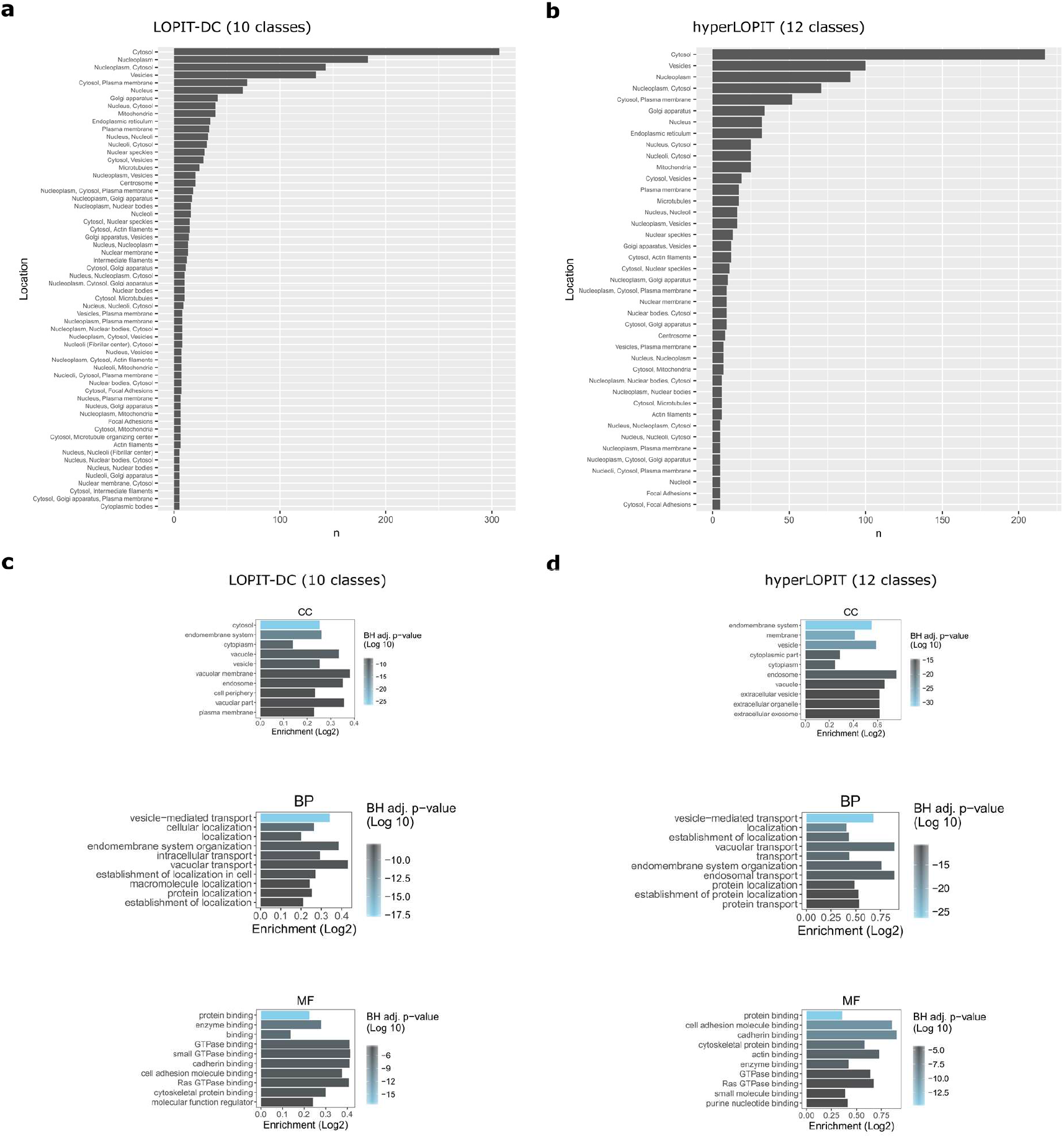
Characterisation of the LOPIT-DC and hyperLOPIT unclassified proteomes. a, b) The subcellular location of the proteins which remained unassigned in the LOPIT-DC and hyperLOPIT data after 5% FDR filtering was investigated in the Cell Atlas database and our barplots show how many of these unlabelled proteins are found as part of the locations listed in the Cell Atlas (the minimum count included in the plots is 5); c, d) The proteins that were not assigned to an organelle in the LOPIT-DC and hyperLOPIT data after 5% FDR filtering were investigated for enrichment with Cellular Component (CC), Biological Process (BP) and Molecular Function (MF) Gene Ontology annotation terms.

In order to gain additional insights into the multilocalising proteome captured by our two distinct workflows we next performed functional Gene Ontology (GO)^74^ term enrichment analysis on the proteins which remained unassigned in both our LOPIT-DC and hyperLOPIT datasets. Regarding GO Biological Process (BP) terms, both the LOPIT-DC- and hyperLOPIT-specific unclassified proteomes are enriched with terms related to vesicle-mediated transport and the endomembrane system (Figures 5c and 5d). Similarly, concerning GO Cellular Component (CC) terms, the unlabelled proteins in both datasets are enriched with terms associated with the cytosol/cytoplasm, endomembrane system, vesicles and endosome (Figure 5c and d). Here, the LOPIT-DC unassigned proteome exhibits an additional overrepresented term related to the plasma membrane and the hyperLOPIT-specific unclassified proteome is enriched with extra terms associated with the extracellular exosome and extracellular vesicles. Importantly, our GO BP and CC term overrepresentation analysis results corroborate the Cell Atlas comparison findings presented in the previous paragraph, according to which the majority of the unlabelled proteins in both of our datasets is annotated as cytosolic in the Cell Atlas and also a large portion of these proteins is assigned to the vesicles and endomembrane system as part of the same database. Moreover, the enrichment of both the LOPIT-DC- and hyperLOPIT-specific unassigned protein pools with general terms referring to macromolecule subcellular localisation might refer to presence of factors which regulate the establishment of protein localisation (or the localisation of molecules other than proteins) within the cell.

Regarding GO Molecular Function (MF) terms, the proteins with an unknown localisation in both the LOPIT-DC and hyperLOPIT data are enriched with terms related to (Ras) GTPase-, cadherin-/cell adhesion molecule-, actin-/cytoskeletal protein- and enzyme-binding (Figures 5c and 5d). Here, the LOPIT-DC-specific unlabelled proteome displays an additional overrepresented term associated with molecular function regulation and the hyperLOPIT unclassified proteins are enriched with an extra term related to purine nucleotide-binding, which possibly refers to transcription factors or RNA-binding proteins. These results provide additional validation and insights on the molecular nature of the multilocalising proteins identified by our two spatial proteomics methods, revealing the presence of protein function regulators as well as interactors of structural components, signalling molecules and nucleic acids among the LOPIT-DC and hyperLOPIT “unknowns”.

### Important biological features can be mapped upon the LOPIT-DC and hyperLOPIT datasets

Aiming to further explore the quality of our data we also examined the clusters present in both the LOPIT-DC and hyperLOPIT datasets in terms of suborganellar resolution. As demonstrated in Figure 6a (and in more detail in Figures S10 and S11), the distributions of several suborganellar structures mapped upon our data exhibit a superb level of agreement between the two datasets. For example, the ER lumen and ER membrane are located at slightly different positions on top of the ER cluster and the ERGIC-cis Golgi is positioned between the ER and GA clusters in both the LOPIT-DC and hyperLOPIT data. Interestingly, these suborganellar structures are better resolved from each other in the LOPIT-DC rather than the hyperLOPIT dataset. Additionally, as expected due to the broad connectivity of the cytoskeleton with most subcellular structures, actin-binding proteins are distributed mainly in the “unknown” area of our plots in both datasets, with some proteins being located close to a variety of organelles. Similarly, our endosomal markers are distributed on top of the plasma membrane and lysosome clusters as well as the unassigned area of the hyperLOPIT PCA plot, while the same proteins exhibit a slightly shifted distribution in the LOPIT-DC data where they are located closer to the ER and in the “unknown” area of the plot. Both distributions are justifiable since the endosome, as part of the endocytic membrane transport pathway, is a very dynamic organelle which recycles between the GA, plasma membrane and lysosomes and is also in contact with the ER^75^.

**Figure 6:**
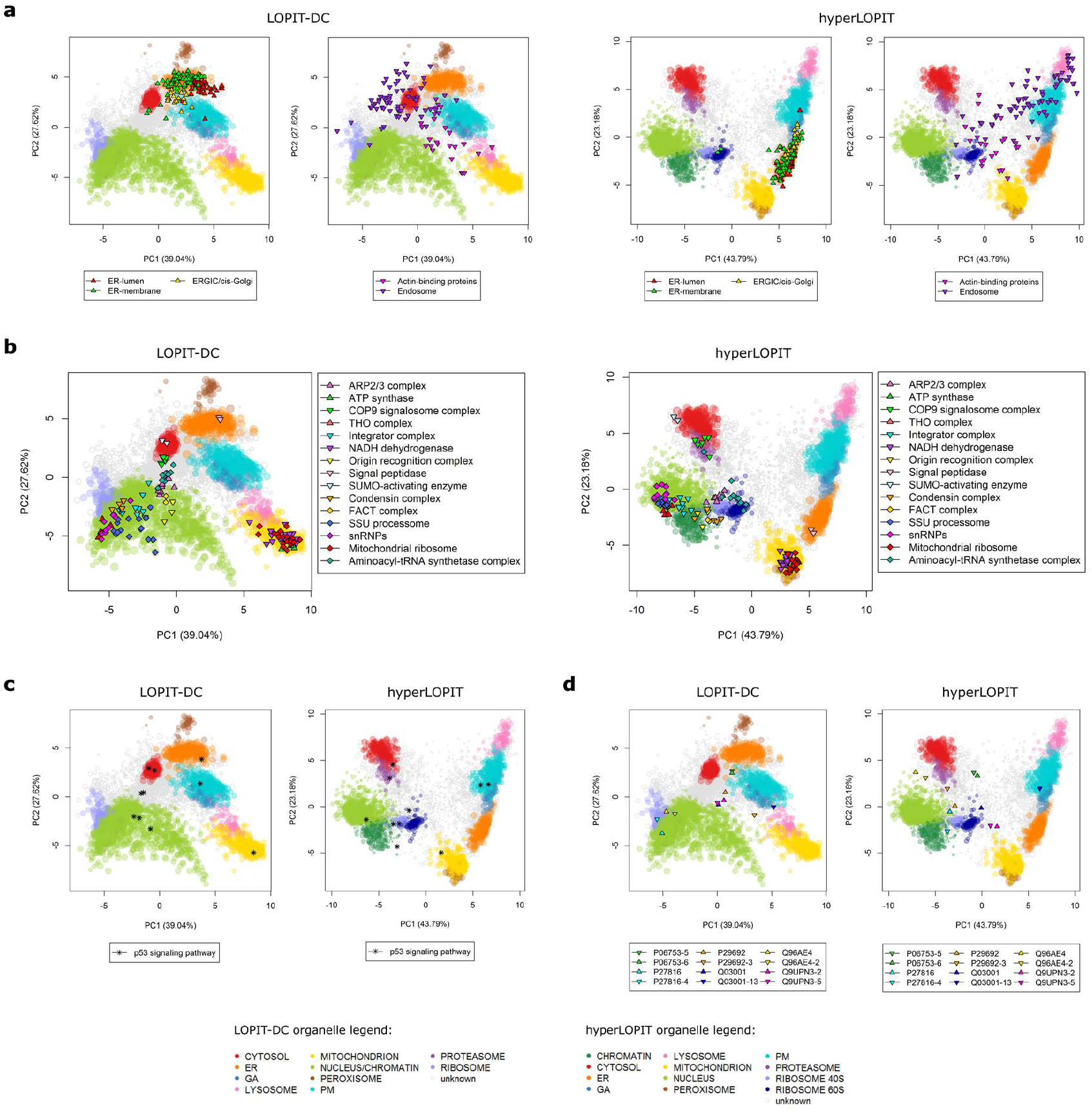
Localisation of suborganellar clusters, large protein complexes, signalling pathways and protein isoforms in the LOPIT-DC and hyperLOPIT datasets. a) Various suborganellar structures plotted upon the LOPIT-DC and hyperLOPIT datasets with assigned proteins; b) Various complexes plotted upon the LOPIT-DC and hyperLOPIT datasets with assigned proteins; c) Proteins involved in p53 signaling plotted upon the LOPIT-DC and hyperLOPIT datasets with assigned proteins; d) Isoform pairs plotted upon the LOPIT-DC and hyperLOPIT datasets with assigned proteins.

We also plotted several large protein complexes upon the PCA plots of the LOPIT-DC and hyperLOPIT data with similar results. As shown in Figure 6b, the majority of complexes we examined exhibit identical distributions in our two datasets. For example, the SUMO-activating enzyme complex and COP9 signalosome are both located within the cytosolic cluster in both the LOPIT-DC and hyperLOPIT data. Similarly, the integrator complex and SSU processome are positioned on top of the nucleus, the NADH dehydrogenase complex and mitochondrial ribosomes are situated within the mitochondrial cluster and the signal peptidase complex is found upon the ER cluster in both of our datasets.

We next sought to investigate the location of distinct components of signalling pathways in our LOPIT-DC and hyperLOPIT datasets. A comprehensive illustration of several pathways which are important for many essential cellular functions is presented in Figures 6c and S12-S21. Interestingly, we observe that the majority of these individual pathway components are found in the same subcellular niche in both datasets. One of the pathways we inspected is the p53 signalling pathway, presented in Figure S12. This pathway plays a crucial role in the control of DNA replication and cell division as well as in cellular responses to different types of stress and has been implicated in many cancers^76^. As exhibited in Figure S12, two components of this pathway can be found in the cytosol, two other of its constituents are classified as PM and ER, one p53 pathway element is situated within the mitochondrion and five additional proteins which belong to this signalling cascade are unassigned in both the LOPIT-DC and hyperLOPIT datasets. Furthermore, since our experiments were performed using the U-2 OS cell line which is a cancer (osteosarcoma) cell line, we also explored pathways which have been found to play critical roles in cancer. For example, we looked at proteins that have been shown to be involved in transcriptional misregulation in cancer, presented in Figure S16. As demonstrated by this figure, twenty-four such proteins can be found in our two datasets. Of these, five are classified to the PM, one is in the GA and fifteen overlap with the nuclear as well as the ribosomal clusters in both the LOPIT-DC and hyperLOPIT datasets. The remaining three components of this pathway visually from our PCA plots are positioned close to the cytosol in the LOPIT-DC dataset and are slightly shifted towards the unassigned area of the plot in the hyperLOPIT data. Importantly, we observed similar results for all the pathways we examined which showcases the outstanding agreement between our two datasets and reflects the functional organisation and networks of the cell. All pathways presented here were plotted according to information available in the KEGG PATHWAY database^77^.

Finally, we also explored the distribution of protein isoforms in our data. As seen in Figure 6d, we could identify six examples where two different isoforms of a protein were present in both of our datasets. In all of these cases, each isoform is mapped to the same subcellular niche in the LOPIT-DC and hyperLOPIT data. For example, both **Q96AE4** isoforms (yellow) were assigned to the same unique subcellular location, the nucleus, in our two datasets. Indeed, this protein is a DNA-binding transcription factor which regulates the expression of the c-Myc gene^78^ and is listed as a nuclear resident in the UniProt and Cell Atlas databases.

In a slightly different case, the Q9UPN3 (pink), P06753 (green), P29692 (orange) and Q03001 (dark blue) isoform pairs presented in the same figure all remained unclassified in both the LOPIT-DC and hyperLOPIT data and therefore overlap with the “unknown” area of our PCA plots, but each isoform of every pair is positioned close to the same organelles in the two datasets. Interestingly, these proteins seem to possess important regulatory roles which could justify their identification as dynamic translocators. In more detail, **Q9UPN3** isoform 2 is a well-studied actin-binding protein which crosslinks actin to other cytoskeletal components, binds to and stabilises microtubules and is involved in the control of focal adhesion assembly and dynamics as well as cell migration and vesicle transport through the trans-Golgi network^79, 80^. This protein also acts as a positive regulator of the

Wnt receptor signalling cascade as it has been shown to be involved in the translocation of the AXIN1/APC/CTNNB1/GSK3B complex from the cytoplasm towards the plasma membrane^81^. According to the UniProt database Q9UPN3 isoform 2 has been found in multiple subcellular locations including the plasma membrane, Golgi apparatus, cytoskeleton and cytoplasm and its localisation is dependent upon its phosphorylation state. Similarly, **P06753** is a tropomyosin chain component, another cytoskeletal element which is listed as a resident of both the cytoskeleton and cytosol in the UniProt and Cell Atlas databases. On the other hand, **P29692** isoform 1 (chosen as the canonical sequence) is an elongation factor which regulates the function of EF-1-alpha and therefore the transfer of aminoacyl-tRNAs to the ribosomes^82^. This protein is described as a resident of the nucleus and cytosol in the UniProt and Cell Atlas databases and indeed, in both of our datasets, it is found between the nuclear and cytosolic clusters and, in the case of the hyperLOPIT data, also close to the two ribosomal subunits which is in agreement with its molecular function. Lastly, **Q03001** and Q03001-13 are positioned on top of the uncharted area and close to the plasma membrane cluster, respectively, in our PCA plots corresponding to both datasets. Similarly to Q9UPN3 and P06753, Q03001 is a dynamic cytoskeletal linker protein which regulates the organisation and stability of intermediate filaments as well as microtubule and actin cytoskeleton networks by acting as an integrator^83^. This protein is also involved in the docking of the dynactin/dynein motor complex to vesicle cargos during retrograde axonal transport. According to UniProt, Q03001 isoform 1 has been found at various cytoskeletal structures throughout the cytoplasm as well as focal contact attachments at the cell membrane and Q03001 isoform 8 has been observed at the plasma membrane, cell cortex and several other cytoskeletal formations.

Finally, **P27816** and P27816-4 (turquoise) were classified to different subcellular compartments to each other in both the LOPIT-DC and hyperLOPIT data. P27816 isoform 1 (the canonical isoform) was assigned to the nucleus while P27816 isoform 4 remained unlabelled in both of our datasets. According to Kitazawa et al.^84^ this protein associates with microtubules and promotes their assembly and according to UniProt it has indeed been identified as part of the cytoskeleton but has also been found in the cytosol, plasma membrane and extracellular or secretory vesicles. In our LOPIT-DC and hyperLOPIT data, P27816 isoform 4 is situated at the border between the nuclear and ribosomal clusters.

## Discussion

The well-established hyperLOPIT workflow allows for the proteome-wide, high-resolution tracking of protein subcellular localisation where multiple organelles are analysed during a single experiment and has been applied to the study of many different biological systems. Since this method provides high-quality, global spatial maps its application can be time-consuming as well as labour- and resource-intensive. Aiming at researchers who do not necessarily seek a maximum resolution-yielding protocol we developed a simpler alternative to hyperLOPIT which we named LOPIT-DC. During the systematic study presented in this manuscript we applied both methods to a human osteosarcoma cell line using identical cell culture and lysis conditions as well as protein identification and quantitation pipelines and data analysis strategies. We compared the results produced via each workflow using a variety of approaches including QSep, a tool which enables the robust quantification of subcellular resolution in spatial proteomics datasets. Our findings indicate that the data generated using hyperLOPIT exhibit the best overall resolution while the dataset obtained using LOPIT-DC closely follows. Our analysis further suggests that the non-chromatin nucleus and chromatin as well as large protein complexes such as the ribosomes and proteasome are not well-separated from each other in our LOPIT-DC data, while the same structures display distinct distributions in the hyperLOPIT dataset. Despite that, the data produced by the two approaches showed excellent agreement regarding protein subcellular localisation prediction which increased even further when the strengths of these methods were integrated using transfer learning. Importantly, both workflows were able to retain crucial information corresponding to suborganellar resolution as well as the localisation of protein complexes and components of important signalling pathways, with extremely high agreement regarding the distribution of individual proteins. Furthermore, the two methods yielded comparable results related to protein isoform-specific subcellular niches with particular isoforms exhibiting very similar distributions in the LOPIT-DC and hyperLOPIT data. Moreover, using information available in the Cell Atlas and Gene Ontology databases we could assign the LOPIT-DC- and hyperLOPIT-specific unlabelled, multilocalising proteomes to subcellular compartments with almost identical results for the two datasets.

In conclusion, both workflows presented in this study can achieve high overall resolution and display excellent reproducibility. The choice regarding which one to use depends on the biological question in mind as well as the amount of starting material, time and resources available. If such matters are of no concern then hyperLOPIT can provide maximum subcellular resolution as the method of choice but in cases of starting material, time or financial constraints the simpler and quicker LOPIT-DC protocol can offer a great all-in-one alternative. As our findings demonstrate, both methods yield reliable, comparable results and can be utilised in the context of dynamic studies or for the mapping of features such as post-translational modifications, protein interactions or isoform behaviour. Importantly, our U-2 OS LOPIT-DC and hyperLOPIT data are the highest-resolution mass spectrometry-based spatial proteomics maps created using human cells to date; these datasets provide a snapshot of the structural organisation of U-2 OS cells and can serve as a reference for future studies on human protein subcellular localisation and its relationship to protein function.

## Materials and Methods

### Cell culture

The U-2 OS human osteosarcoma cell line was a generous gift from Professor Emma Lundberg (SciLifeLab Stockholm and School of Biotechnology, KTH). The cells were grown at 37 °C and 5% CO2 in McCoy’s 5A medium (Sigma) supplemented with sodium bicarbonate, 10% foetal bovine serum (Biosera) and 1% GlutaMax™ (Life Technologies), without antibiotics.

### Sample preparation

Samples were prepared as described in^85^ and^72^. U-2 OS cells were trypsinised, washed and resuspended in a gentle lysis buffer (0.25 M sucrose, 10 mM HEPES pH 7.4, 2 mM EDTA, 2 mM magnesium acetate, protease inhibitors). They were then lysed using a ball-bearing homogeniser and spun at 200 x g, 5 min, 4 °C to remove unlysed cells.

In parallel, a chromatin extraction step was performed using approximately 5-15 million cells (10-20% of the total number of cells used per experiment) according to^85^.

### hyperLOPIT subcellular fractionation

Samples for hyperLOPIT were treated with nuclease and then fractionated using an iodixanol density gradient as described in^85^ and^72^. Briefly, a cell lysate from approximately 280 million cells per average experiment was first separated into a cytosol-enriched and a crude membrane fraction using 6% and 25% (w/v) iodixanol-containing solutions and centrifugation at 100,000 x g, 90 min, 4 °C. The supernatant was stored and the membrane fractions situated at the interface of the iodixanol layers collected and centrifuged to get rid of any residual cytosolic contamination. The samples were then resuspended in 25% (w/v) iodixanol, underlaid beneath a linear gradient of 8%, 12%, 16% and 18% (w/v) iodixanol solutions and fractionated by centrifugation at 100,000 x g, 8 h, 4 °C. After ultracentrifugation approximately 20-22 fractions were collected, pelleted several times at 100,000 x g, 1 h, 4°C to wash away the iodixanol and stored at −80 °C.

The cytosol-enriched supernatant was precipitated with five volumes of cold acetone overnight at – 20 °C. The obtained precipitated pellet and membrane pellets were resolubilised in 8 M urea, 0.2% SDS and 50 mM HEPES pH 8.5. Protein concentration was measured using the BCA protein assay kit (Thermo Fisher Scientific) according to the manufacturer’s instructions.

60-70 ug of protein per fraction were reduced with TCEP, alkylated with MMTS, digested with trypsin and labelled with isobaric tagging reagents as previously described^85^. For our hyperLOPIT samples, each tag from a TMT10plex kit (Thermo Fisher Scientific) was split in half (essentially making the labelling scheme a 20plex) and used to label all the membrane fractions (some were pooled to ensure adequate protein amounts) as well as the cytosol- and chromatin-enriched samples. Three TMT10plexes were used to label three biological replicates.

After labelling, peptides were pooled into 10plexes, cleaned with C18 SepPak cartridges and fractionated using high-pH reverse phase chromatography. The resulting fractions corresponding to each TMT10plex set were orthogonally combined into 18-22 samples for downstream MS analysis.

### LOPIT-DC subcellular fractionation

Samples for LOPIT-DC were fractionated using differential centrifugation (Table 1). Besides the cell lysis and data analysis steps, the method differs from Itzhak *et al.*^67^ in the extended centrifugation scheme and, most importantly, in its ability to capture all subcellular niches in a single experiment. A cell lysate from approximately 70 million cells per average experiment was separated into 10 fractions using the Eppendorf 5804 R for the first centrifugation step and the Optima™ MAX-XP Beckman benchtop ultracentrifuge with the TLA-55 rotor for the rest. All pellets and the last supernatant were stored at −80 °C.

The final supernatant was precipitated with five volumes of cold acetone overnight at −20 °C. The obtained precipitated pellet and membrane pellets were resolubilized in 8 M urea, 0.15% SDS and 50 mM HEPES pH 8.5. Protein concentration was measured using the BCA protein assay according to the manufacturer’s instructions.

50 ug of protein per fraction were reduced, alkylated, digested and TMT-labelled as previously described^85^. One TMT10plex kit was used to label all the membrane and cytosol-enriched fractions in three biological replicates. The TMT11-131C tag was used to label the chromatin-enriched fraction.

After labelling, peptides were pooled into 10plexes, cleaned with C18 SepPak cartridges and fractionated using high-pH reverse phase chromatography. The 11^th^ tag was added to each 10plex just before RP-HPLC. The resulting fractions corresponding to each TMT10plex set were orthogonally combined into 18 samples for downstream MS analysis.

### SDS-PAGE and immunoblotting

To make sure that subcellular fractionation was conducted successfully in both cases, SDS-PAGE^86^ and western blotting were performed using the antibodies from^85^. Proteins were separated on Mini-PROTEAN TGX Precast Gels (Bio-Rad) and transferred to nitrocellulose or polyvinylidene fluoride (PVDF) membranes using the Trans-Blot Turbo Transfer System (Bio-Rad). Signal was detected using the ECL Prime Western Blotting Detection Reagent (GE Healthcare) kit according to the manufacturer’s instructions.

### SPS (Synchronous Precursor Selection)-MS^3^ on the Orbitrap Fusion Lumos

All mass spectrometry runs were performed on an Orbitrap Fusion™ Lumos™ Tribrid™ instrument coupled to a Dionex Ultimate™ 3000 RSLCnano system (Thermo Fisher Scientific) with parameters from^85^.

### Raw data processing and quantification

Raw files were processed with Proteome Discoverer v1.4 (Thermo Fisher Scientific) using the Mascot server v2.3.02 (Matrix Science). The SwissProt sequence database for Homo *sapiens* (canonical and isoform, 42,118 sequences, downloaded on 04/11/2016) was used along with common contaminants from the common Repository of Adventitious Proteins (cRAP) v1.0 (48 sequences, adapted from the Global Proteome Machine repository). Precursor and fragment mass tolerances were set to 10 ppm and 0.6 Da, respectively. Trypsin was set as the enzyme of choice and a maximum of 2 missed cleavages were allowed. Static modifications were: methylthio (C), TMT6plex (N-term) and TMT6plex (K). Dynamic modifications were: oxidation (M) and deamidated (NQ). Percolator was used to assess the false discovery rate (FDR) and only high confidence peptides were retained. Additional data reduction filters were: peptide rank = 1 and ion score > 20.

Quantification at the MS^3^ level was performed within the Proteome Discoverer workflow using the centroid sum method and an integration tolerance of 2 mmu. Isotope impurity correction factors were applied. Each raw peptide-spectrum match (PSM) reporter intensity was then divided by the sum of all intensities for that PSM (sum normalisation). Protein grouping was carried out according to the minimum parsimony principle and the median of all sum-normalised PSM ratios belonging to each protein group was calculated as the protein group quantitation value. Only proteins with a full reporter ion series were retained. Finally, proteins identified as cRAP were removed for downstream analysis.

### Machine learning and multivariate data analysis

#### A) SVM-based prediction of protein localisation

Data analysis was performed using the R^87^ Bioconductor^88^ packages MSnbase^89^ and pRoloc^44^ as described in^69^. Briefly, 579 manually curated marker proteins were used to define 12 subcellular locations: cytosol, proteasome, nucleus, chromatin, 40S ribosome, 60S ribosome, peroxisome, mitochondrion, lysosome, Golgi apparatus, plasma membrane and endoplasmic reticulum (supplemental quantitation table). These constitute our “core organelle markers”, proteins known to localise to one specific subcellular niche. Supervised machine learning using a support vector machine (SVM) classifier with a radial basis function kernel was employed in order to predict the localisation of unlabelled proteins. In the case of the LOPIT-DC data classification was performed using both 12 and 10 marker classes: in the latter case the pairs nucleus/chromatin and ribosome 40S/ribosome 60S were merged to form single classes. Following the protocol in^85^, one hundred rounds of fivefold cross-validation was employed (creating five stratified test/train partitions) to estimate algorithmic performance. This protocol features an additional round of cross-validation on each training partition to optimise the free parameters of the SVM, sigma and cost, via a grid search. Based on the best F1 score (the harmonic mean of precision and recall), for the hyperLOPIT dataset the best sigma and cost were 0.01 and 8, respectively. The best sigma and cost for the LOPIT-DC data annotated with 12 marker classes were 0.1 and 16 and for the dataset with 10 classes they were 0. 01 and 16. All proteins assigned to a specific subcellular niche by SVM-based classification were ordered according to their SVM scores and a threshold was set to achieve a 5% FDR based on agreement with the UniProt and Gene Ontology databases.

#### B) Data integration by transfer learning

To show the complementary nature of the hyperLOPIT and LOPIT-DC methods at predicting subcellular location we applied a transfer learning (TL) algorithm^71^. The TL method allows one to integrate heterogeneous datasets (a primary and an auxiliary dataset) for optimal classification. Following the protocol described in^71^, the hyperLOPIT dataset was used as the primary source and the LOPIT-DC as the auxiliary source. Labelled marker proteins common in both datasets were extracted and the hyperLOPIT and LOPIT-DC quantitative protein profiles were used as input to the k-nearest neighbor transfer learning (knntl) algorithm. Three different experiments were conducted: (1) using the hyperLOPIT data only, (2) using the LOPIT-DC data only and finally (3) using both hyperLOPIT and LOPIT-DC data. As per the SVM classifier, one hundred rounds of fivefold crossvalidation were used to estimate the optimal number of nearest neighbours for the k-nearest neighbour (k-NN) classifier. These were 5 and 5 for the hyperLOPIT and LOPIT-DC datasets, respectively. In the k-NN transfer learning framework we also need to estimate the parameter theta which is a vector of weights (one per organelle) used to control the amount of primary (hyperLOPIT) and auxiliary (LOPIT-DC) data to use in classification. We tested all weight combinations of 0, 0.5 and 1 for each organelle class. The median theta weight vector was picked over 100 rounds of crossvalidation based on the highest F1 score. The optimal theta weight for integrating the datasets was theta = (0, 0.75, 0.5, 0.5, 1, 1, 0.5, 0.5, 0.5, 0.5, 1, 1) for the cytosol, ER, GA, lysosome, mitochondrion, nucleus, chromatin, peroxisome, PM, proteasome, 40S ribosome and 60S ribosome, respectively.

#### C) QSep analysis

The QSep function which is freely available as part of the pRoloc package^44^ was used to quantify the resolution of the LOPIT-DC and hyperLOPIT datasets (for more details see^73^). QSep calculates cluster separation by comparing the average Euclidean distances within and between subcellular clusters. These distances always refer to one specific organelle marker cluster and the distances within clusters are usually smaller than the ones between clusters, except in cases of overlapping subcellular niches. To enable reliable comparison of such distances within a single experiment but also across different studies QSep further divides each value by the reference within-cluster average distance, as follows:

QSep calculates all between- and within-cluster average distances. These distances are then divided column-wise by the respective within-cluster average distance. For example, for a dataset with only two spatial clusters, *C*_1_ and *C*_2_, we would obtain

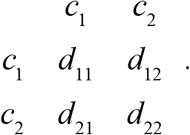

Normalised distances here represent the ratio of between-to-within average distances, i.e. how much larger the average distance between clusters *C_i_*. and *C_j_*. is compared to the average distance within cluster *C_i_*.

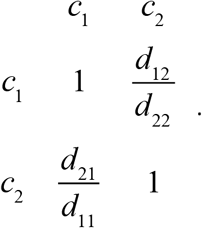

Following this calculation the normalised distance matrix ceases being symmetric and the normalised distance ratios are proportional to the tightness of the reference cluster (along the columns).

As demonstrated in the above example, the resulting distance value is informative of how much the average distance between two clusters is greater than the average distance within a cluster, the reference within-cluster distance here being a measure of how compact a cluster is. The resolution metric used by QSep is not influenced by the number of classes used for its computation and performs consistently well when provided with different organelle marker annotation. However, subcellular marker definition does affect the resolution assessment scoring with low quality marker lists yielding suboptimal results^73^. Resolution measurements acquired via the QSep function can be visualised using quantitative cluster separation heatmaps and boxplots (see Figures 3a and 3b and^73^).

## Data availability

All protein level datasets are available in the R Bioconductor pRolocdata package (Gatto, L. & Breckels, L. M. (2018). pRolocdata: Data accompanying the pRoloc package. R package version 1. 18.0, https://github.com/lgatto/pRolocdata, https://bioconductor.org/packages/pRolocdata). Further information should be directed to the lead contact, K.S.L.

## Author Contributions

A.G., N.K.B. and K.S.L. designed, planned and performed the experiments and data analysis. T.S.S. and L.G. performed data analysis. L.M.B. provided practical guidance regarding data analysis. A.G., N.K.B. and K.S.L. wrote the manuscript with contributions from L.M.B. and L.G. Figures were prepared by N.K.B., A.G., T.S.S. and L.G. C.M.M. and O.M.C. advised on the content and layout of the manuscript.

## Acknowledgments

A.G. was funded through the Alexander S. Onassis Public Benefit Foundation, the Foundation for Education and European Culture (IPEP), the A. G. Leventis Foundation and the Embiricos Trust Scholarship of Jesus College Cambridge. T.S.S. and L.G. were supported by a Wellcome Trust Grant (grant no. 110170/Z/15/Z). L.M.B. and C.M.M. were supported by a Wellcome Trust Technology Development Grant (grant no. 108467/Z/15/Z). O.M.C. is a Wellcome Trust Mathematical Genomics and Medicine student supported financially by the School of Clinical Medicine, University of Cambridge. K.S.L. is a Wellcome Trust Joint Senior Investigator (grant no. 110170/Z/15/Z). We would like to thank Dr Mike Deery for performing the mass spectrometry for this project.

